# Para- and transcellular transport kinetics of nanoparticles across lymphatic endothelial cells

**DOI:** 10.1101/2023.04.12.536598

**Authors:** Jacob McCright, Jenny Yarmovsky, Katharina Maisel

**Affiliations:** Department of Bioengineering, University of Maryland College Park, College Park, MD, USA

**Keywords:** lymphatic endothelial cells (LECs), surface chemistry, micropinocytosis, macropinocytosis, endocytosis, immunotherapy

## Abstract

Lymphatic vessels have received significant attention as drug delivery targets, as they shuttle materials from peripheral tissues to the lymph nodes, where adaptive immunity is formed. Delivery of immune modulatory materials to the lymph nodes via lymphatic vessels has been shown to enhance their efficacy and also improve bioavailability of drugs when delivered to intestinal lymphatic vessels. In this study we generated a three-compartment model of a lymphatic vessel with a set of kinematic differential equations to describe the transport of nanoparticles from surrounding tissues into lymphatic vessels. We used previously published data and collected additional experimental parameters, including transport efficiency of nanoparticles over time, and also examined how nanoparticle formulation affected the cellular transport mechanisms using small molecule inhibitors. This experimental data was incorporated into a system of kinematic differential equations and non-linear, least squares curve fitting algorithms were employed to extrapolate transport coefficients within our model. The subsequent computational framework produced some of the first parameters to describe transport kinetics across lymphatic endothelial cells and allows for the quantitative analysis of the driving mechanisms of transport into lymphatic vessels. Our model indicates that transcellular mechanisms, such as micro- and macropinocytosis, drive transport into lymphatics. This information is crucial to further design strategies that will modulate lymphatic transport for drug delivery, particularly in diseases like lymphedema, where normal lymphatic functions are impaired.

## Introduction

Nanoparticle-based drug delivery has received significant attention in past decades, culminating in the recent COVID-19 lipid nanoparticle-based RNA vaccines. Nanoparticles provide several advantages to free drug formulations:they can increase drug stability [1] and load [2], be targeted to specific tissues or cells [3], and facilitate sustained release of drugs from the nanoparticle core [4]. To enter the relevant cellular compartments, nanoparticles are required to be transported across various biological barriers, including cell monolayers at the intestinal epithelium or blood and lymphatic endothelium, for example. Nanoparticle transport across these cellular barriers occurs via paracellular or transcellular transport mechanisms. Transcellular transport mechanisms include micropinocytosis, macropinocytosis, and/or exocytosis on both sides of the cell layer (**Figure 1A**), often with concentration gradients driving the dominant direction of the transport. Nanoparticle transport across cellular barriers can be modeled using systems of differential equations utilizing kinetic theories also applied to pharmacokinetic models [5] A complex system of equations can be used to define the contributions of the individual transport mechanism to the overall transport of nanoparticles (**Equations 1-3**). This system of equations can be applied for transport across any system of single cell layers or other three-compartment models.

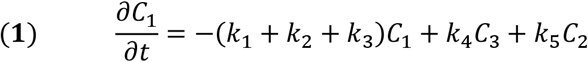

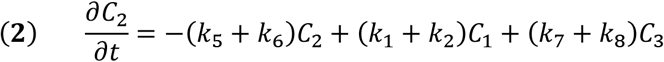

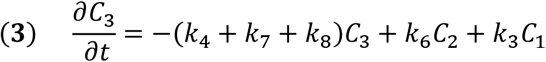

Lymphatic vessel-targeted drug delivery has received significant attention in recent years, largely owing to the fact that lymphatic vessels transport nanoparticle-sized materials from the peripheral tissue to the draining lymph nodes, where the adaptive immune response is shaped. Delivering therapeutics that modulate the immune response directly to the lymph nodes has been shown to enhance their efficacy. Nanoparticles 10-250 nm in size have been shown to be preferentially transported by lymphatic vessels and will accumulate effectively in the lymph nodes. Recent studies have shown that both transcellular and paracellular mechanisms are key in nanoparticle entry into lymphatic vessels [6-8]. Research from our group has shown that the specific transport mechanisms used by lymphatic endothelial cells are dependent on nanoparticle surface chemistry, specifically PEG density [8]. We found that coating 100 nm nanoparticles with hydrophilic, neutrally charged poly(ethylene glycol) (PEG) leads to nanoparticle transport via both micropinocytosis and paracellular transport. Using computational models to study the kinetics behind these findings can aid in our understanding of how nanoparticle surface chemistry affects transcellular processes [9-14] [15]. Here, we sought to model the kinetics of nanoparticle transport across lymphatic vessels. In this paper, we derive the equations and rate coefficients describing transport kinetics of nanoparticles, taking into account both transcellular and paracellular mechanisms. Additionally, we use transport data collected experimentally and simplify **Equations 1-3** into a four-part problem of endocytosis, exocytosis, and paracellular transport, with the assumption that transport is driven in the direction of the concentration gradient from the interstitial tissue into the lumen of the vessel (**Figure 1B, Equations 4-6**). The resulting mathematical and computational framework can be used to extrapolate transport kinetics across similar cell layer problems and could be integrated with more complex machine learning-based techniques, like artificial neural networks, to predict the contribution of different transport mechanisms.

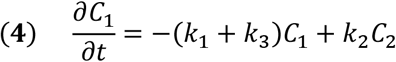

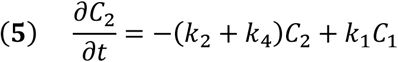

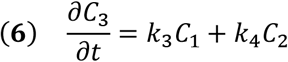

**Figure 1:**
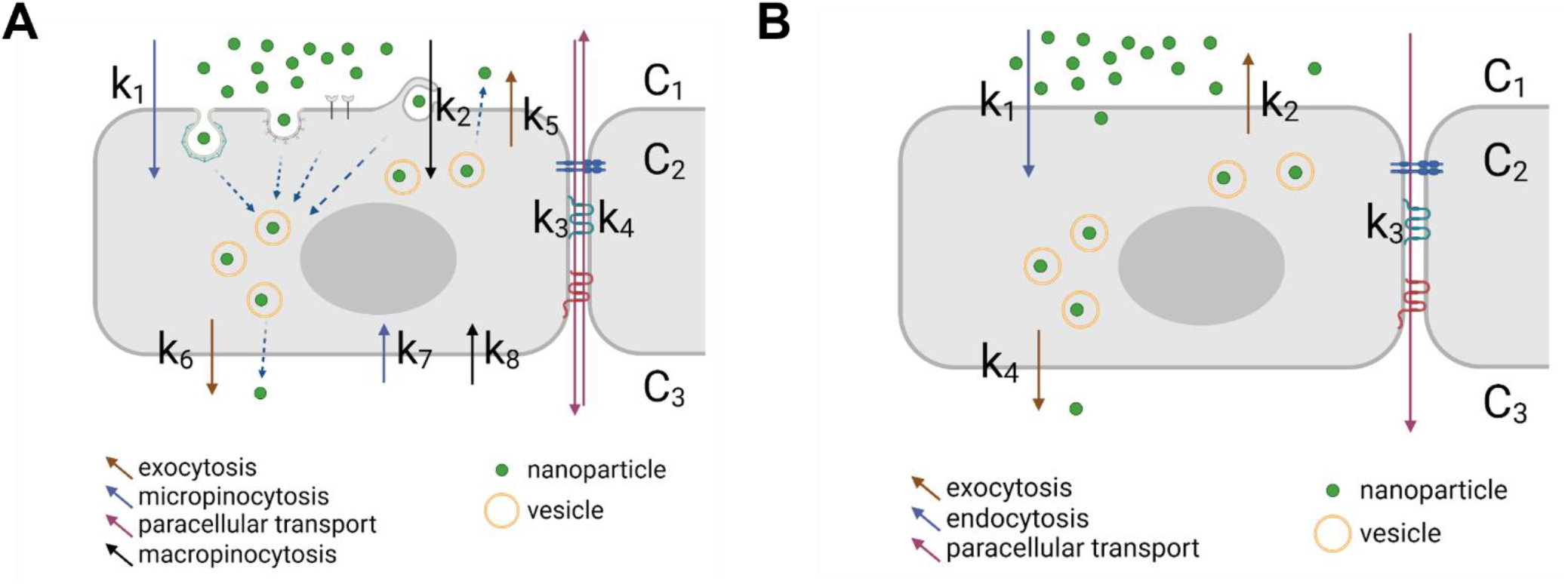
Cellular mechanisms used to transport nanoparticles across cell barriers such as epithelial surfaces and vessel walls. **A)** complex transport considering all potential variables and **B)** simplified transport considering only endocytosis, paracellular transport, and exocytosis. Concentrations depict each compartment with C_1_ = nanoparticle-rich compartment, C_2_ = intracellular compartment, C_3_ = nanoparticle-poor compartment. Full arrows and associated k values represent the kinetics of exocytosis (brown), micropinocytosis (blue), macropinocytosis (black), and paracellular transport (pink). Figure created using bioRender.

## Methods

### Nanoparticle Formulation

Fluorescent carboxyl (COOH)-modified polystyrene (PS) nanoparticles (Thermo Fisher Scientific) were covalently modified with 5 kDa MW methoxy-PEG-amine (NH_2_) (Creative PEGworks), as previously described [16]. Nanoparticles with different PEG conformations were generated using previously described methods [17]. Briefly, PS-COOH particles were suspended at 0.1% w/v in 200 mM borate buffer (pH = 8.2). PEG was conjugated to nanoparticles using 7 mM N-Hydroxysulfosuccinimide (NHS) (Sigma) and 0.02 mM 1-Ethyl-3-(3-dimethylaminopropyl) carbodiimide (EDC) (Invitrogen). The reaction was performed on a rotary incubator at room temperature for at least 4 hours. Nanoparticles were collected using 100k MWCO centrifugal filters (Amicon Ultra; Millipore) and washed with deionized (DI) water. Nanoparticles were resuspended at 1% w/v in DI water and stored at 4°C.

### Nanoparticle Characterization

Dynamic light scattering (DLS) was used to measure the hydrodynamic diameter and polydispersity index (PDI) of nanoparticles. Phase analysis light scattering (PALS) was used for measuring ζ-potential (NanoBrook Omni). Measurements were performed using a scattering angle of 90° at 25°C. Measurements were based on intensity of reflected light from scattered particles.

### PEG Density Characterization

PEG density was determined using a previously published method [17, 18]. Briefly, 5kDa PEG-NH2 (Creative PEGworks) conjugated to fluorescein isothiocyanate (FITC) was conjugated to fluorescent (AlexaFluor®555) 100 nm carboxyl-modified nanoparticles (Creative PEGworks). Using FITC-PEG-NH_2_ standards, PEG grafting distance (D) and PEG density were estimated using the Flory radius of PEG (Rf) based off of fluorescence intensity. The Flory radius of a polymer chain is defined as Rf ∼ αN^3/5^, where N is the degree of polymerization, and α is the effective monomer length. An unconstrained 5 kDa PEG chain has an Rf of 5.4 nm and occupies 22.7 nm^2^. PEG density and conformation can be correlated to the ratio of Rf/D, with Rf/D < 1-1.5 yielding a mushroom, 1-1.5< Rf/D > 4 yielding a brush, and Rf/D > 4 yielding a dense brush conformation.

### Nanoparticle Transport

LEC permeability was assessed using an established in vitro model that recapitulates in vivo lymphatic transport [19]. Briefly, primary human LECs (hLECs, Promocell C-12217) were seeded on 1.0 µm pore size, 12 mm transwell inserts (Falcon) at 200,000 cells/cm2 and cultured in EGM2 (PromoCell) at 37°C and 5% CO2 for 48 h. Cells were pre-treated with 1 µm/s transmural flow to simulate the interstitial fluid flow experienced in the tissue microenvironment. hLECs were treated with 10 ug/mL nanoparticles in the top compartment and both top and bottom compartment were sampled every 1 h for up to 24 h. Fluorescence intensity was measured using a plate reader (Tecan) and nanoparticles transported was calculated using a standard curve. Transport experiments were performed in EGM2 without growth factors to avoid confounding effects of growth factors on transport mechanisms. To probe transport mechanism, the following transport inhibitors were used: 100 nM Adrenomedullin (Abcam ab276417), 62.5 µM Dynasore (Sigma D7693), or 62.5 µM Amiloride (Sigma A7410). Transport inhibitors were applied 2 hours prior to introduction of nanoparticles. LEC monolayer integrity was confirmed after experiments using immunofluorescence.

### Computational Model Solving

Equations 1-3 were generated to model the three-compartment model of nanoparticle transport across LECs (**Figure 1A**). The following assumptions were made to simplify the model 1) Transport between cells is unidirectional along the concentration gradient since [C1] >> [C3] (k4 = 0) and 2) Reuptake of nanoparticles from compartment C3 to C2 is negligible (k7, k8 = 0). Under these assumptions, equations 1-3 become equations 4-6 (**Figure 1B**).

Experimental data of C1 and C3 were used to extrapolate C2, assuming that nanoparticle mass and fluorescence was conserved. This data was normalized to C = C/Ctotal. The normalized concentration over time data was used to estimate k parameters using MatLab ‘lsqcurvefit’ function. Levenberg–Marquardt, or the damped least-squares method, was used to solve the nonlinear least squares optimization problem:

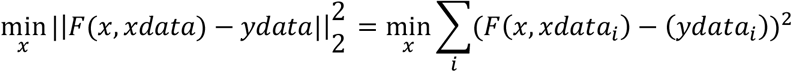

For parameter estimation, k values were constrained to be positive to match the kinetics outlined in equations 4-6 and **Figure 1B**. In the presence of Adrenomedullin, parameter k3 was set to zero, as the addition of Adrenomedullin prevents paracellular transport.

## Results

### Transport Efficiency of Nanoparticles Across LECs can be Fitted to a Three-Compartment Kinetic Model

To be able to fully model the transport kinetics within the three compartments (**Figure 2A)**, C1 (top compartment, interstitial tissue where nanoparticles are injected), C2 (intracellular), and C3 (bottom compartment, vessel lumen that leads to the lymph nodes), we first collected data on concentration change over time within C1 and C3 (**Figure 2B**). We formulated 40-150 nm PEGylated nanoparticles [16, 17] from unmodified 103 ± 5 nm and 40 ± 2 nm. Addition of PEG increased nanoparticle diameter to 122 ± 6 nm (partial PEG) and 142 ± 3 nm (dense PEG), and 43 ± 2 nm (partial PEG) and 49 ± 3 nm (dense PEG) (**Supplementary Figure 1A**). Pegylated nanoparticles also displayed a near-neutral ζ – potential (**Supplementary Figure 1B**). Using these nanoparticles, we found that C1 was reduced to 60-70% of the initial nanoparticle concentration placed in the top well, after which C1 increased again when nanoparticles were exocytosed into the top compartment (**Figure 2B**). We also confirmed that densely PEGylated nanoparticles were transported the most efficiently compared to partially PEGylated nanoparticles, with 5.3 ± 0.2% nanoparticles transported into C3 for densely PEGylated 100 nm nanoparticles, compared to 4.1 ± 0.2% for partially PEGylated nanoparticles (**Figure 2B**). Similarly, 40 nm densely PEGylated nanoparticles had 7.9 ± 0.5% of nanoparticles transported into C3, compared to 5.5 ± 0.2% for the partially PEGylated nanoparticles (**Figure 2B**).

**Figure 2:**
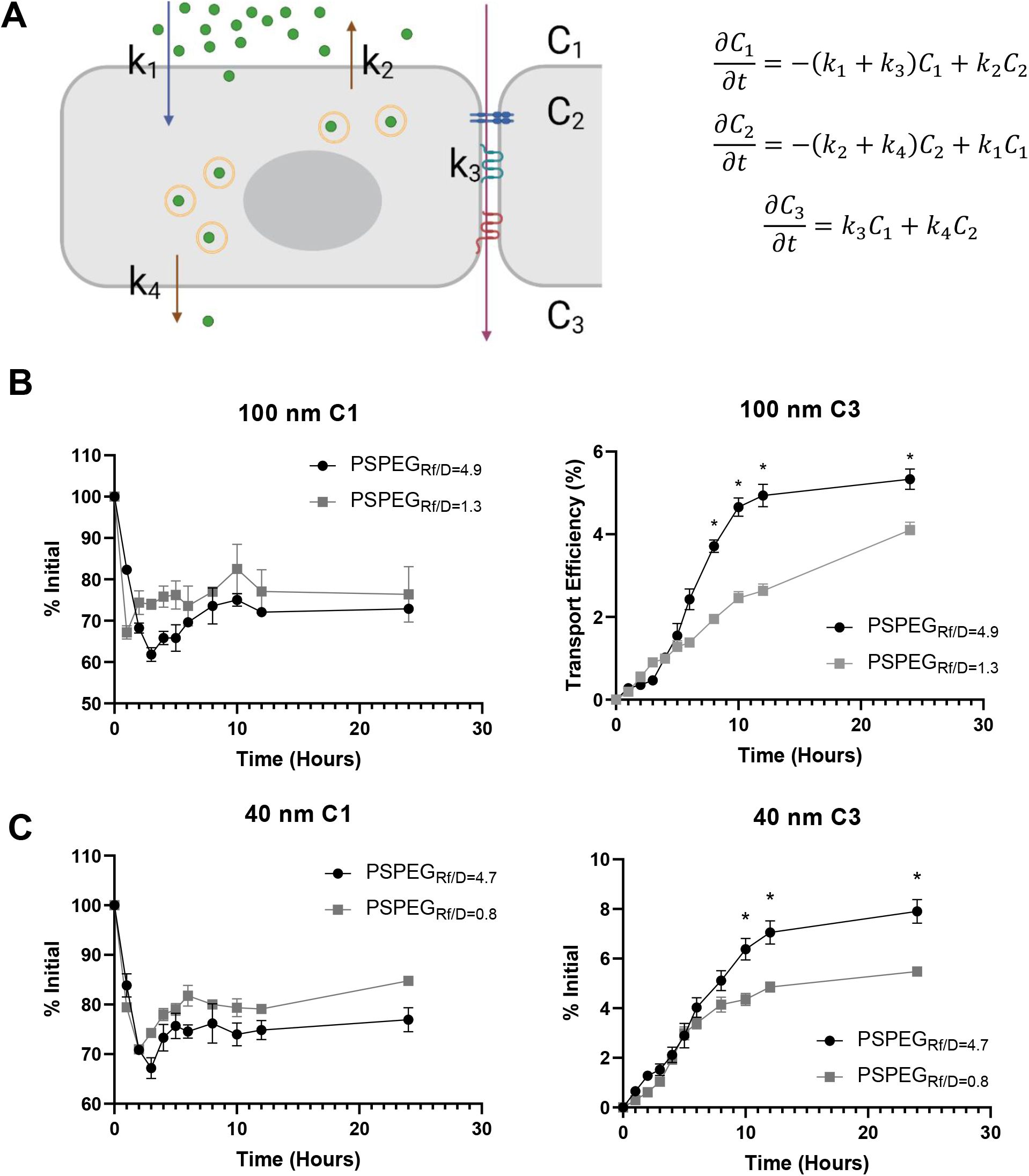
PEG Coating Improves Transport of 100 and 40 nm NP Across LECs. **A)** Schematic of the transport model and associated differential equations. **B)** % 100 nm nanoparticle transport (densely PEGylated, PSPEG_Rf/D=4.9_, low density PEGylated, PSPEG_Rf/D=1.3_) in the apical (C1, left) and basolateral (C3, right) compartments. **C)** % of 40 nm nanoparticle transport (densely PEGylated, PSPEG_Rf/D=4.7_, low density PEGylated, PSPEG_Rf/D=0.8_) in the apical (left) and basolateral (bottom) compartments. Data presented as mean ± SEM (*p<0.05) and representative of n ≥ 3 repeat experiments. Figure created using bioRender.

We then used C1 and C3 data (**Figure 2B)** and a non-linear, least-squares, curve fitting algorithm (**Figure 3**) to estimate the kinetics (k) parameters in the system of differential equations (**Table 1**). Based on these k values, we can see that the initial uptake and release of nanoparticle at the top compartment interface (k1 and k2) is the dominant reaction, indicated by the order of magnitude difference compared to k3 and k4. Another trend to note is that k3, describing paracellular transport, is the smallest parameter. This indicates that cellular mechanisms, including endocytosis and micropinocytosis are driving nanoparticle transport across LEC barriers. Importantly, transport trends with respect to formulation are captured within the model. k4, a key parameter for measuring the transport of nanoparticles into the simulated vessel, increases from 0.13 µg mL^-1^ hr^-1^ for the larger 100 nm PSPEG_Rf/D=4.9_ nanoparticles to 0.26 µg mL^-1^ hr^-1^ for the smaller 40 nm PSPEG_Rf/D=4.7_ nanoparticles.

**Table 1:** Calculated K values of system of differential equations describing nanoparticle transport across LECs for different nanoparticle formulations.

| Rate Constants ($\mu\text{g mL}^{-1} \text{ hr}^{-1}$) | 100 nm PSPEG<sub>Rf/D=4.9</sub> | 100 nm PSPEG<sub>Rf/D=1.3</sub> | 40 nm PSPEG<sub>Rf/D=4.7</sub> | 40 nm PSPEG<sub>Rf/D=0.8</sub> |
| --- | --- | --- | --- | --- |
| k1 | 1.9 | 1.5 | 1.5 | 1.2 |
| k2 | 3.9 | 3.5 | 4.0 | 3.2 |
| k3 | 0.013 | 0.012 | 0.014 | 0.011 |
| k4 | 0.13 | 0.074 | 0.26 | 0.21 |

**Figure 3:**
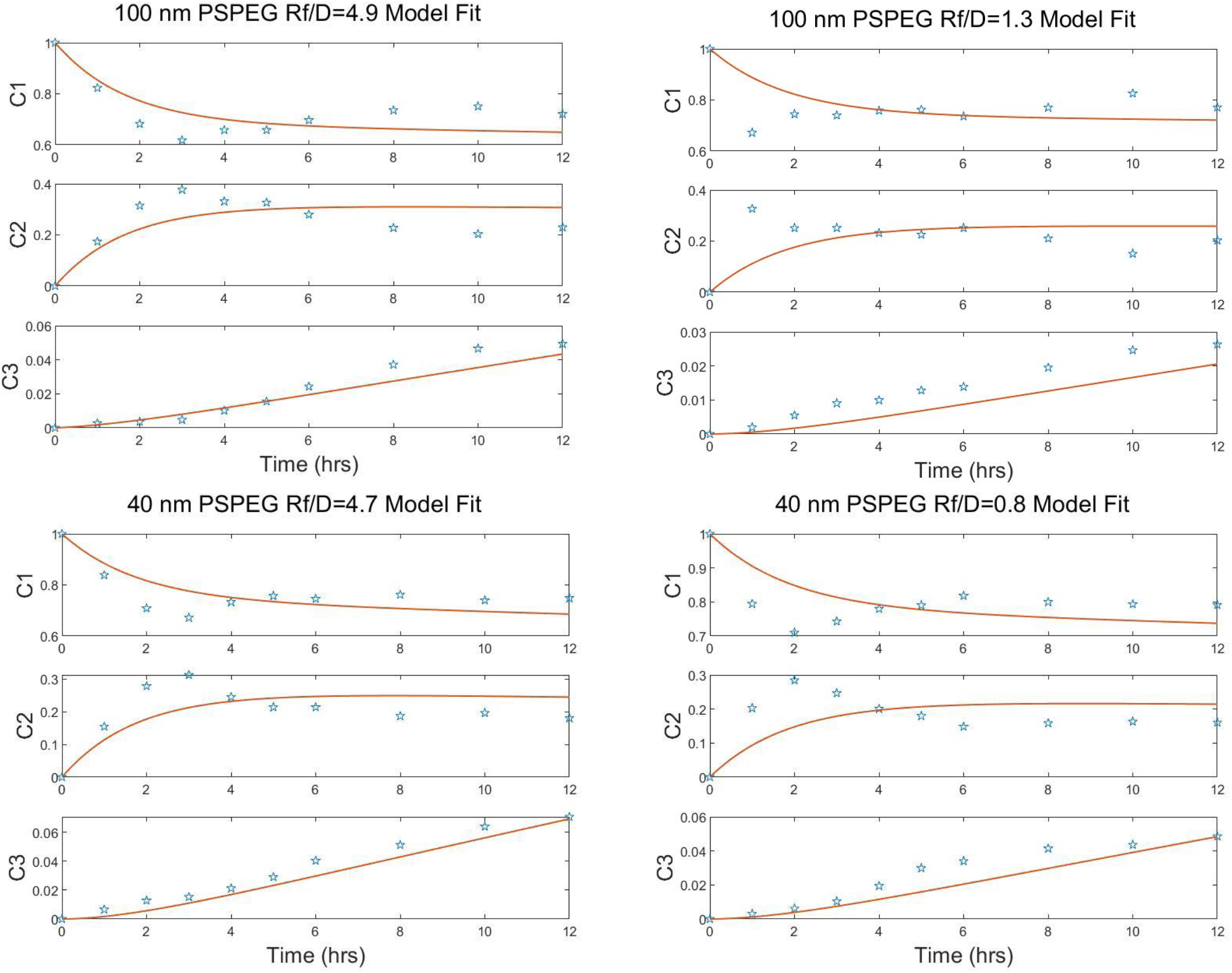
Nanoparticle transport across LECs modeled with a 3-compartment kinetic model. Normalized experimental concentration data (stars) fitted against system of differential equations (solid line).

### Size and surface chemistry of nanoparticles affect their transport via macropinocytosis across LECs

Next, we investigated how the kinetics changed for different cellular transport mechanisms. Again, we used partially and densely PEGylated, 40 – 150 nm nanoparticles. As in our prior study, we found that 100 nm densely PEGylated nanoparticles were not transported by macropinocytosis but that both paracellular and transcellular transport were involved, indicated by reduced transport upon introduction of transport inhibitors (**Figure 4A**). Interestingly, for partially PEGylated nanoparticles, all transport mechanisms were involved in nanoparticle accumulation in C3 (**Figure 4A**). The introduction of transport inhibitors generally reduced the depletion of nanoparticles from C1: for both 100 nm PSPEG_Rf/D=4.9_ and PSPEG_Rf/D=1.3,_ inhibiting micropinocytosis reduced this phenomenon the greatest, albeit not significantly different (**Figure 4B**).

**Figure 4:**
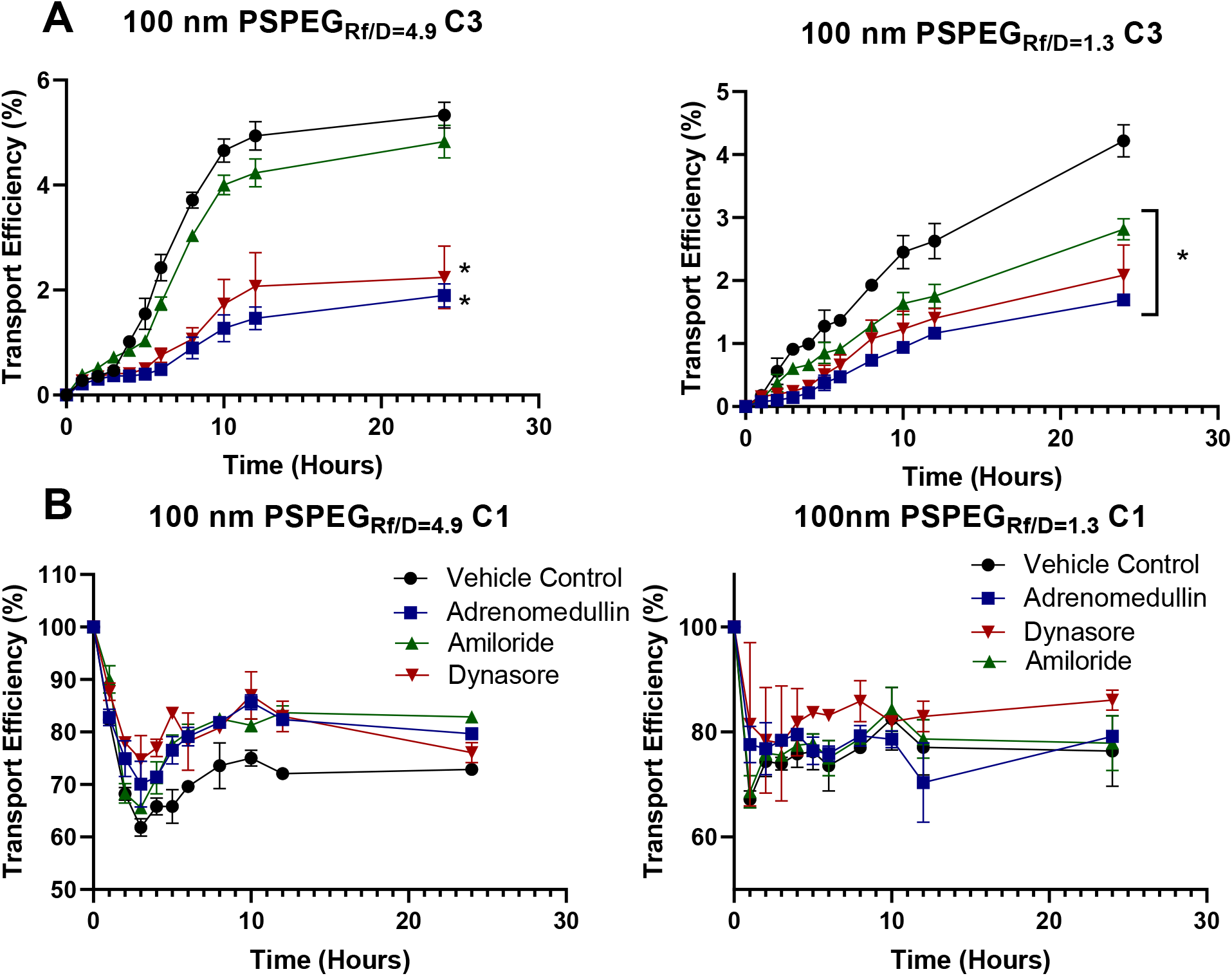
Nanoparticle transport across LECs is mediated by micropinocytosis, macropinocytosis, and paracellular transport. Percent 100 nm nanoparticle transport (densely PEGylated, PSPEG_Rf/D=4.9_, low density PEGylated, PSPEG_Rf/D=1.3_) in the **A)** basolateral (C3) and **B)** apical (C1) compartments in the presence of transport inhibitors. Data presented as mean ± SEM (*p<0.05) and representative of n ≥ 2 experiments.

**Figure 5:**
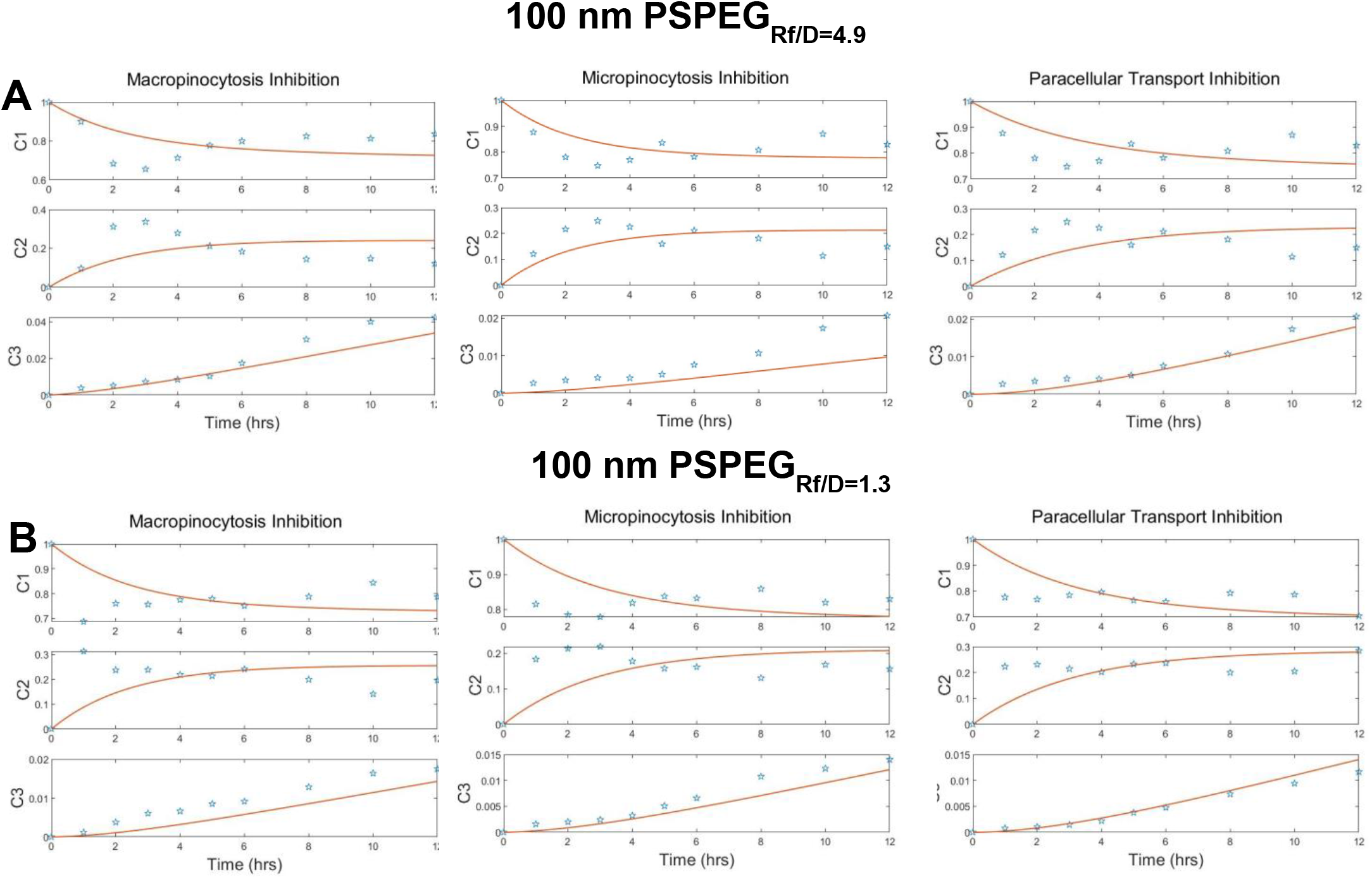
Kinetic model captures transport mechanisms governing nanoparticle transport across LECs. Percent 100 nm nanoparticle transport of **A)** densely PEGylated, PSPEG_Rf/D=4.9_ and **B)** low density PEGylated, PSPEG_Rf/D=1.3_) with inhibition of micropinocytosis, macropinocytosis, or paracellular transport. Normalized experimental concentration data (stars) fitted against system of differential equations (solid line) including different transport inhibitors.

This transport data including transport inhibitors was then incorporated into our three-compartment kinetic model and fitted to the system of differential equations using a non-linear, least-squares, curve fitting algorithm, resulting in curve fitting (**Figure 4C-D**). Solved parameters k for equations with respect to nanoparticle formulation can be found in **Tables 2-3**. From these k values, we can see trends in the presence of inhibitors similar to the control transport experiments: For both densely and partially PEGylated nanoparticles, k1 and k2 (uptake and re-release of nanoparticles into C1) are larger than k3 and k4 (transcellular and paracellular transport of nanoparticles into C3).

**Table 2:** Calculated k values of system of differential equations describing transport of 100 nm PSPEG across LECs in the presence of different transport inhibitors.

| Rate Constants<br>($\mu\text{g mL}^{-1} \text{ hr}^{-1}$) | 100 nm PSPEG<sub>Rf/D=4.9</sub> | +Amiloride | +Dynasore | +Adrenomedullin |
| --- | --- | --- | --- | --- |
| k1 | 1.9 | 1.2 | 1.0 | 1.70 |
| k2 | 3.9 | 3.0 | 3.6 | 2.2 |
| k3 | 0.013 | 0.02 | 0.2 | 0 |
| k4 | 0.13 | 0.10 | 0.041 | 0.088 |

**Table 3:** Calculated k values of system of differential equations describing transport of 100 nm PSPEG across LECs in the presence of different transport inhibitors.

| Rate Constants<br>($\mu\text{g mL}^{-1} \text{ hr}^{-1}$) | 100 nm PSPEG<sub>Rf/D=1.3</sub> | +Amiloride | +Dynasore | +Adrenomedullin |
| --- | --- | --- | --- | --- |
| k1 | 1.5 | 1.0 | 0.7 | 1.90 |
| k2 | 3.5 | 2.8 | 2.6 | 2.2 |
| k3 | 0.012 | 0.01 | 0.18 | 0 |
| k4 | 0.074 | 0.10 | 0.057 | 0.055 |

Our computational model captures how the inclusion of transport inhibitors affects mechanisms used for transport. Removing paracellular transport resulted in an increase in k1 for partially PEGylated nanoparticles. When micropinocytosis was inhibited, k1 dropped from 1.9 to 1.0 µg mL^-1^ hr^-1^ and 1.5 to 0.7 µg mL^-1^ hr^-1^ for densely and partially PEGylated 100 nm nanoparticles, respectively. k3, which represents paracellular transport, in contrast increased nearly 10-fold, suggesting that an increased concentration gradient drives more nanoparticles across LECs via paracellular transport. Even though our experimental data suggests that inhibition of macropinocytosis does not significantly modulate densely PEGylated, 100nm nanoparticle transport across LECs, our computational data shows a reduction in k1 upon treatment with amiloride from 1.9 to 1.2, suggestive of the complex relationship between the regulation of different transcellular transport mechanisms that exist in biological systems.

## Discussion

In this study we probed how nanoparticle formulation parameters, including size and surface chemistry, influenced both the transport efficiency and transport mechanism into lymphatic vessels. We found that increasing the density of PEG on the surface of nanoparticles improved transport efficiency, with maximal transport efficiency occurring with the smaller, 40 nm densely PEGylated nanoparticles. We also observed that nanoparticle transport mechanism is dependent on formulation, notably that macropinocytosis does not drive transport for the 100 nm fully PEGylated nanoparticles as it does for other formulations. Using these experimental results, we fitted our data into a set of differential equations describing the three-compartment problem including endocytosis, exocytosis, and paracellular transport. This computational framework produced parameters to describe transport kinetics and allow for the quantitative analysis of driving mechanisms of transport and formulation parameters.

Lymphatic vessels exist throughout the entire body and are known for transporting cells, fluid, and particulates from peripheral tissues to the local draining lymph nodes, where the adaptive immune response is formed. Lymphatics are important drug delivery targets as they transport immune modulatory therapies to the lymph nodes and improve vaccine and immunotherapy efficacy [20]. A study from Triacca et al provided some of the preliminary evidence that LECs themselves serve as significant barriers todelivery into lymphatic vessels [19]. Developing the in vitro lymphatic transport model that is also used in this study, they demonstrated that when micropinocytosis was inhibited with Dynasore, transport efficiency of albumin and dextran across the LECs decreased significantly – a clear indication that LECs serve as barriers. We also recently demonstrated first insights on the transport mechanisms involved in the transport of nanoparticles across lymphatic barriers. We found that both paracellular and transcellular transport mechanisms were key in crossing lymphatic barriers, with LECs relying on clathrin-mediated endocytosis to mediate transport of PEGylated nanoparticles [17]. Our current study is an extension of this work, demonstrating that macropinocytosis, in particular, is affected by both size and surface chemistry. Others have demonstrated that macro- and micropinocytosis are involved in nanoparticle transport across endothelial cells in tumors and the blood brain barrier [21-27]. Rabanel et al demonstrated that nanoparticles coated with 5 kDa PEG were taken up primarily via macropinocytosis pathways in brain endothelial cells [28]. Tehrani et al found that inhibiting micropinocytosis reduced transcytosis across brain endothelial cells by 60% for 5 kDa PEG-coated nanoparticles, while transcytosis of 2 kDa PEG-coated nanoparticles was reduced only by 25% after inhibiting micropinocytosis [29]. These findings suggest that endocytosis drives nanoparticle uptake and that transcytosis pathways may differ with nanoparticle size and different amounts and density of PEG, corroborating the results in this study. Additionally, studies have demonstrated that nanoparticle transport across brain microvascular endothelium can be enhanced by taking advantage of existing receptor-mediated transcytosis, such as that of albumin (clathrin/caveolin-dependent) [25, 30-33].

Computational methods and frameworks to model drug delivery into tissues and across endothelial barriers have been used to study the regulation of nanoparticle transport and to guide nanoparticle design to target tissues of interest. One of the key considerations when applying a computational model to physiological phenomena is deciding what type of model to employ. In our study, we generated a three-compartment model of the lymphatic endothelium and used the Levenberg–Marquardt algorithm for parameter estimation to solve our non-linear least squares parameter optimization problem for our system of ODEs. This algorithm was used as it is more robust than the more common Gauss-Newton algorithm, especially when the data is not well-behaved or the starting parameters are far from the solution parameters, as is often the case when modeling physiological data. [34] Saqlain et al endeavored to model levodopa concentration in the brains of Alzheimer’s patients using a classical system of ODEs and a novel approach using stochastic differential equations[35]. They found that the stochastic model better fit the physiological data. This shift to stochastic modeling has correlated with the emergence of neural network-based and artificial intelligence-based modeling for drug delivery[36]. Lu et al employed a novel neural-ODE system to predict T-DM1 conjugate concentration in patients based on patient characteristics (age, sex, race, height, weight, region) and dosing regimen[37]. They validated the implementation of the neural-ODE system against lightGBM and LSTM methods for predicting pharmacokinetics and found that their neural-ODE system better recapitulated the clinical data compared to traditional models. Cattaneo and Zunino generated a computational model based on fundamental filtration and transport to model the interplay between blood perfusion, fluid exchange with the interstitial volume, mass transport in the capillary bed, and transport through the capillary walls and into the surrounding tissue at the microscale level[38]. In their work, they solved for the transport of small molecule drugs and nanoparticles from capillaries into surrounding tumor spaces. Using this model, they were able to quantitatively demonstrate that using nanoparticle intermediaries for drug delivery was an optimal delivery strategy compared to bolus injection of free drug in the tumor site. Groh et al also employed a computational model to examine drug transport into tumors [14]. In this study they simulated solid tumors, and through the solving of their computational model, were able to identify key pharmacokinetic parameters that govern how far model drugs penetrate model tumors.

In addition to computational modeling of drug delivery on the larger tissue level, studies have also sought to model drug and nanoparticle transport across, and interactions with, cells and cell layers. Wei-Chun Chou et al developed a pharmacokinetic model to examine gold nanoparticle kinetics in rat models. Through the development of their model, they examined how size affected kinetics[39]. Interestingly, based on their physiological data, they were able to modify a classical pharmacokinetic modeling framework to hypothesize nanoparticle-specific pathways and kinetics with respect to size. While their model focused on uptake and clearance of nanoparticles at the organ scale, formation of their model indicated that classical approaches employed for small molecule pharmacokinetic modeling did not translate to nanoparticle modeling and that administration route-specific data and modeling is needed to improve approaches to modeling nanoparticle pharmacokinetics. A key consideration from their study is that the lymphatics and lymph nodes were not included in their model, highlighting the gap in knowledge regarding nanoparticle pharmacokinetics in these key tissues. A recent paper from Khan et al employed a similar transwell-based model as outlined in our study [15]. They generated a three-compartment model and fitted observed transport data to a system of kinematic equations to solve for the transport coefficients governing their model. This study highlighted how artificial neural networks (ANNs) can be used as a method to statistically model and solve kinematic transport equations similar to the ones set forth in our study. In their model, they observed a steady accumulation of nanoparticle within the intracellular compartment, in contrast to the rapid uptake and release observed in our model. Collectively, these studies demonstrate the utility of computational models and how they can be employed to better understand how the transport of key agents in tissues of interest is regulated. Our work builds on these concepts by using computational methods to examine what mechanisms are driving transport, as well as using these methods to quantitatively describe how formulation parameters can affect transport efficiency.

One of the interesting findings from our study is the nanoparticle uptake and release back into the top well. This phenomenon has been observed in other studies, including one by Georgieva et al, that showed how caveolin-mediated uptake of nanoparticles by endothelial cells peaked at 30 min before release back into the environment [40]. This relatively rapid uptake and release of nanoparticles is similar to what occurred in our study, reinforcing that endocytosis pathways are key when examining nanoparticle transport. Another study from Fiorentino et al observed a similar phenomenon where 20-100 nm PS nanoparticles were rapidly taken up by blood outgrowth endothelial cells and released back into cell culture within four hours. Furthermore, they observed that nanoparticles localized with caveolin and that inhibiting endocytotic pathways with chemical inhibitors, including Dynasore, prevented nanoparticle uptake [41]. This endocytosis-mediated uptake and translocation of nanoparticles is similarly observed in our study, with the administration of the micropinocytosis inhibitor Dynasore decreasing the rapid uptake of nanoparticles seen in untreated controls. Rapid uptake and release of nanoparticles has been observed in a variety of cell types, including epithelial cells, fibroblasts, and macrophages [42-45].

## Conclusion

In summary, our study demonstrates that a dense coating of PEG (Rf/D > 4) is required to maximize transport across lymphatic barriers. These findings are consistent with prior work demonstrating that PEG enhances uptake and transport across other endothelial barriers, as well as the cellular mechanisms involved in this transport. This work is one of the first to closely examine the kinetics of nanoparticle transport across the lymphatic barrier. Our computational framework can be integrated with more complex machine learning-based techniques, such as artificial neural networks, to predict the contributions of different transport mechanisms and drive formulation strategies for nanoparticles that will maximize transport across lymphatics and to the LNs, which is crucial for future development of immune modulatory therapeutic strategies.

## Abbreviations

(PEG): poly(ethylene glycol)
(PS): polystyrene
(DLS): dynamic light scattering
(PDI): polydispersity index
(PALS): phase analysis light scattering
(R_f_): Flory radius
(D): grafting distance
(LECs): lymphatic endothelial cells
(ODE): ordinary differential equation
(NHS): N-Hydroxysulfosuccinimide
(3- dimethylaminopropyl): 1-Ethyl-3-
(EDC): carbodiimide
(ANNs): artificial neural networks

## Acknowledgements

We would like to thank the UMD Bioworkshop core facility and Michele Kaluzienski for manuscript editing. This work was funded by NIH MIRA R35 (KM), NSF CAREER Award (KM), the AHA Predoctoral Research Fellowship (JM), and Clark mid-career doctoral fellowship (JM).

